# Using intracerebral microdialysis to probe the efficacy of repurposed drugs in Alzheimer’s disease pathology

**DOI:** 10.1101/2022.01.14.476357

**Authors:** Christiana Bjorkli, Mary Hemler, Joshua B. Julian, Axel Sandvig, Ioanna Sandvig

**Affiliations:** Department of Neuromedicine and Movement Science, Faculty of Medicine and Health Sciences, Norwegian University of Science and Technology, Trondheim, Norway; Department of Neurology, St. Olav’s Hospital, Trondheim, Norway; Princeton Neuroscience Institute, Princeton University, Princeton, NJ, USA; Department of Clinical Science, Faculty of Neuroscience, Umeå University, Sweden

## Abstract

All disease-targeting drug trials completed to date have fallen short of meeting the clinical endpoint of significantly slowing cognitive decline in Alzheimer’s disease patients. Even the recently approved drug Aducanumab, has proven effective in removing amyloid-β, but does not reduce cognitive decline. This emphasizes the urgent need for novel therapeutic approaches that could reduce several AD neuropathologies simultaneously, eventually leading to improved cognitive performance. To validate whether our mouse model replicates AD neuropathology as observed in patients, we characterized the 3xTg AD mouse model to avoid premature translation of successful results. In this study we have repurposed two FDA-approved drugs, Fasudil and Lonafarnib, targeting the Wnt signaling and endosomal-lysosomal pathway respectively, to test their potential to attenuate AD pathology. Using intracerebral microdialysis, we simultaneously infused these disease-targeting drugs between 1-2 weeks, separately and also in combination, while collecting cerebrospinal fluid. We found that Fasudil reduces intracellular amyloid-β in young, and amyloid plaques in old animals, and overall cerebrospinal fluid amyloid-β. Lonafarnib reduces tau neuropathology and cerebrospinal fluid tau biomarkers in young and old animals. Co-infusion of both drugs was more effective in reducing intracellular amyloid-β than either drug alone, and appeared to improve contextual memory performance. However, an unexpected finding was that Lonafarnib treatment increased amyloid plaque size, suggesting that activating the endosomal-lysosomal system may inadvertently increase amyloid-β pathology if administered too late in the AD continuum. Taken together, these findings lend support to the application of repurposed drugs to attenuate AD neuropathology at various therapeutic time windows.

**One Sentence Summary:** Here we circumvented the blood-brain barrier for drug delivery aimed at attenuating AD neuropathology.

## INTRODUCTION

Alzheimer’s disease (AD) is the leading cause of dementia in older adults. Symptoms feature progressive neurodegeneration, followed by impairments in memory, cognitive and visuospatial function (*1*). Brain pathology in AD patients is characterized by abiotrophic neuronal cell death, vascular abnormalities, extracellular deposits of amyloid-β (Aβ), and neurofibrillary tangles (NFTs) (*1*). In most cases, the cause of AD is not known. However, rare inherited versions of AD are often caused by mutations that result in increased production of Aβ (*2, 3*) and lead to familial forms of the disease. Both amyloid plaques and NFTs are believed to interfere with cytoskeletal integrity as well as disrupting axonal transport, and the healthy function of synapses and neurons (*4*). Research within the field has primarily focused on late-stage disease pathology and a central reason for recent failures in clinical trials in AD may thus be the timing of intervention, as most clinical trials begin at advanced stages of the disease, after irreversible neuronal death has already occurred (*5, 6*). It is therefore our view that research efforts need to also focus on earlier stages of AD, when therapeutic interventions would likely be more effective.

Cerebrospinal fluid (CSF) levels of Aβ and tau protein demonstrate high accuracy for identifying incipient AD as well as predicting the development and rate of cognitive decline in the clinic (*4, 7-9*). Lower concentrations of CSF Aβ42 and higher concentrations of total tau (t-tau) protein have been used to distinguish AD patients from cognitively normal age-matched controls and to predict the transition from mild cognitive impairment (MCI) to AD (*4, 7-9*). Studies indicate that abnormal levels of CSF Aβ manifest several years prior to the appearance of subjective memory complaints, making CSF Aβ the current earliest biomarker for AD (*10, 11*). In the early phases of AD, there is evidence that CSF Aβ42 levels significantly increase, initially reflecting the rate of Aβ42 production in the brain (*12*). However, as the disease progresses, AD patients have diminished CSF Aβ42 levels due to a reduction of its clearance and the formation of amyloid plaques in the brain parenchyma (*13*). Recent studies support the hypothesis that the accumulation of Aβ peptides within the brain arises from an imbalance in the production and clearance of Aβ, and the ability to clear Aβ diminishes with age (*14*). Levels of CSF tau serve as an indication of its pathogenesis in the cerebral cortex; both t- and phosphorylated (p)-tau levels increase during progression of AD and correlate with cognitive decline (*15*). It is believed that the decrease in CSF Aβ42 reflects its aggregation and deposition in the brain parenchyma, whereas the increase in CSF tau reflects its extracellular release after neuronal degeneration and NFT formation (*9*). Thus, these biomarkers are considered to be directly linked to the molecular pathogenesis of AD. Since memory impairment manifests in many other diseases in addition to AD, biomarkers are often left as the sole criteria for accurate diagnosis.

Over the past 25 years, there has been an abundance of research on the development of therapies aimed at delaying or halting the progression of AD (*16, 17*). Despite this research, disease-targeting drugs have repeatedly failed to demonstrate success in clinical trials (*18*). Aβ immunotherapy in mouse models of AD has been shown to reduce intracellular Aβ and amyloid plaque accumulation and to lead to a greater clearance of p-tau pathology; however, aggregates of hyperphosphorylated tau remained unaffected by Aβ antibody immunization treatment (*19*). This suggests that Aβ targeting therapies may be more effective at the preclinical stages of AD, but once cognitive decline appears in tandem with tau pathology, tau-targeting drugs may be necessary (*20*). Moreover, the most commonly used AD animal models are mice that overexpress human genes associated with familial AD that result in the formation of amyloid plaques. However, AD is defined by the presence and interplay of both amyloid plaques and NFT pathology (*21*). There has thus been a poor track record of success in AD clinical trials, and this is in part due to the premature translation of successful results in animal models that mirror only limited aspects of AD pathology in patients. Here we have used the 3xTg AD mouse model (*22*) which harbors human AD-related genetic mutations resulting in amyloid plaques and NFTs.

In this study we aimed to target the blood-CSF barrier, formed by the epithelium of the choroid plexuses, for delivery of drugs attenuating AD pathology in the brain parenchyma. First, we repurposed Fasudil, which is used to treat cerebral vasospasm. Fasudil works by targeting Rho-associated protein kinase (ROCK) in the Wnt-PCP signaling pathway (**Fig. 1**), effectively derailing the synaptotoxic cascade of Aβ production. Previous research has demonstrated that ROCK kinases can induce the processing of amyloid precursor protein (APP) to the toxic Aβ42 peptide and that this can be prevented by ROCK inhibition (*23*). Wnt-PCP synaptic signaling is triggered from the increased presence of Aβ protein, and increased expression of Dickkopf WNT Signaling Pathway Inhibitor 1 (Dkk1) has been shown in post-mortem AD brains and in animal models of Aβ pathology (*24, 25*). Moreover, activating autophagy (**Fig. 2**) has increasingly been considered as a potential therapeutic treatment for AD, with animal and cellular models showing that autophagy activators can reduce the levels of misfolded and aggregation proteins, prevent the spread of tau, and reduce neuronal loss (*26-28*). The second drug we repurposed is Lonafarnib, originally a cancer drug and an autophagic activator, which has been found to direct abnormal tau protein into lysosomes, and thereby remove tau before it can form NFTs (*26*).

**Figure 1.**
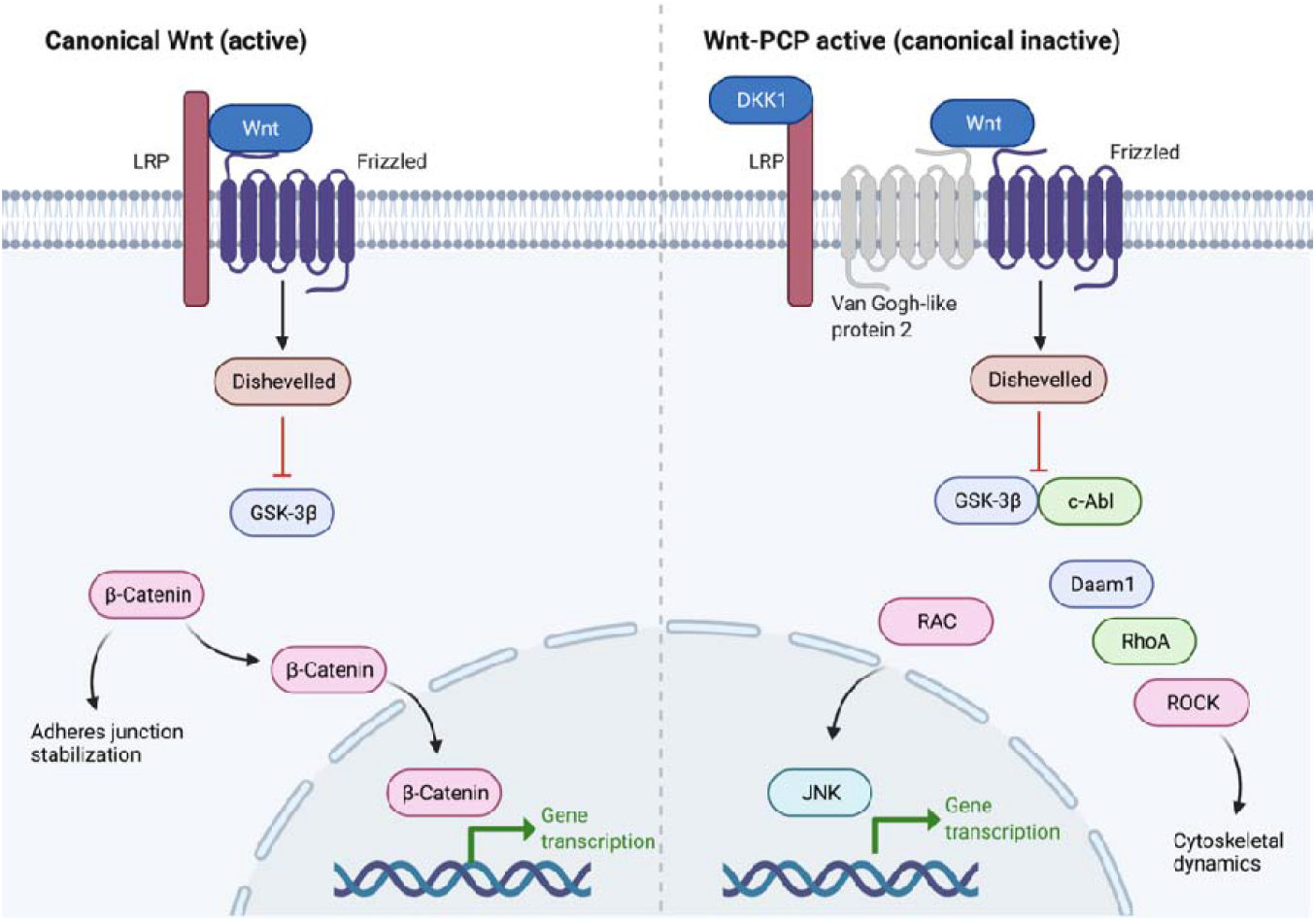
Schematic of the canonical Wnt and Wnt-PCP pathways. Aβ has been shown to activate the Wnt-PCP pathway through the ability of Aβ to induce Dkk1. Dkk1 then prevents the binding interaction between LRP6 and frizzled, activating Wnt-PCP signaling and blocking canonical Wnt-β-catenin activity. In the Wnt-PCP pathway, the two arms diverge below disheveled, acting via Daam/RhoA/ROCK to regulate cytoskeletal dynamics and JNK/c-Jun to regulate gene transcription. Figured adapted from (*24*). Abbreviations; Daam1: disheveled associated activator of morphogenesis 1; Dkk1: Dickkopf-1; GSK-3β: glycogen synthase kinase-3β; JNK: c-Jun N-terminal kinase; LRP: low-density lipoprotein receptor-related protein; PCP: planar cell polarity; RhoA: Ras homolog family member A; ROCK: Rho-associated coiled-coil containing protein kinase; Wnt: Wingless-related integration site. Figure created with biorender.com.

**Figure 2.**
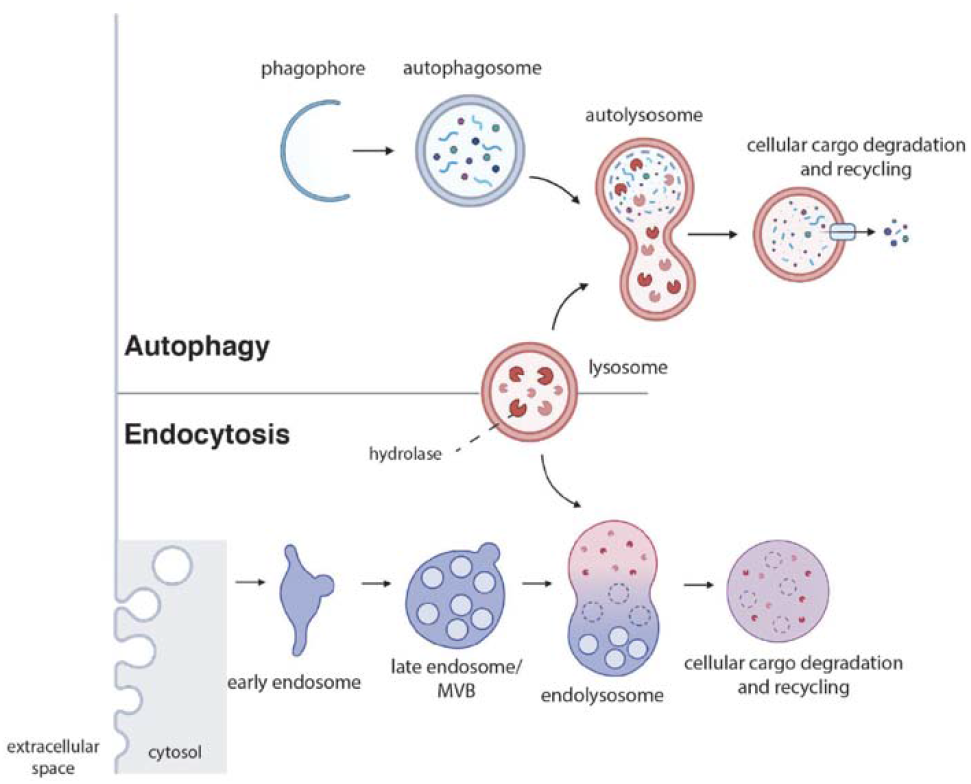
Overview of the autophagic and endocytic pathways. In macroautophagy, the phagophore encloses a portion of cytoplasm that contains organelles and large proteins destined for degradation, forming an autophagosome. The autophagosome then fuses with a lysosome, forming an autolysosome. Lysosomal hydrolases then degrade the internalized material and recycle components back to the plasma membrane. Endocytosis first involves internalization of extracellular material and plasma membrane, which are then delivered to the early endosome. After progressive maturation of the early endosome into a late endosome (also referred to as a multivesicular body), the late endosome fuses with a lysosome. As in autophagy, cellular cargo in the lysosome is then degraded via acid hydrolases. Therefore, both the autophagic and the endocytic pathways converge on the lysosome. Figure adapted from (*31*) and created with biorender.com.

Current AD drug development is limited by the blood-brain barrier (BBB), as more than 98% of all molecular drugs do not penetrate it. Although several lines of research have been developed to investigate transport functions across the BBB, *in vivo* brain microdialysis seems to be one of the most suitable techniques for characterizing the influx and efflux transport functions across the barrier under physiological and pathological conditions (*29*). We predicted that drugs infused into the lateral ventricle f 3xTg AD mice would reach deep cerebral structures, such as the hippocampus (HPC), and that such chronic drug delivery would reduce CSF sink action towards drugs (*30*). We additionally hypothesized that the earlier the biochemical events leading to the deposition of amyloid plaques and NFTs were targeted, the more effective treatment would be and that drug treatment could rescue contextual memory deficits in mice.

## RESULTS

### The 3xTg AD mouse model displays Alzheimer’s disease-related neuropathology

To validate whether our mouse model replicated AD neuropathology as observed in patients, and to assess genetic drift in our own colony (*32*), we first characterized the 3xTg AD mouse model. In the present characterization study intracellular Aβ immunoreactivity is observed in frontal and sensory areas, the HPC, and parts of the cerebellum as early as 1 month of age (**Fig. 3A, B**). Regarding extracellular amyloid plaque accumulation, these are observed first at 13 months of age in the dorsal subiculum (dSub; **Fig. 3A, B**). The observed amyloid plaques are immunoreactive to both McSA1 (Aβ38-42) and OC (fibrillar Aβ) antibodies, suggesting that these plaques consist of fibrillar Aβ38-42. In addition, reactive microglia (Iba1 and TREM2 immunolabelling) surround deposited plaques in this mouse model (**Fig. 3C, D**). At 1 month of age, there is Aβ38-42 immunoreactivity in large parts of the cortex, but not Aβ42 immunoreactivity. Oligomeric Aβ is not present at 1 month of age (**Supplementary Fig. 1**), whereas fibrillar Aβ (OC) is present in the frontal cortex. Moreover, amyloid plaques immunoreactive to Aβ38-42 are first apparent at 13 months of age in dSub, whereas plaques immunoreactive to Aβ42 are first apparent at 15 months of age in dSub (**Fig. 3B**). MC1 immunoreactivity (AD-specific conformation modification of tau) is found within OC-positive amyloid plaques in the dSub of 3xTg AD mice (**Fig. 3E**).

**Figure 3.**
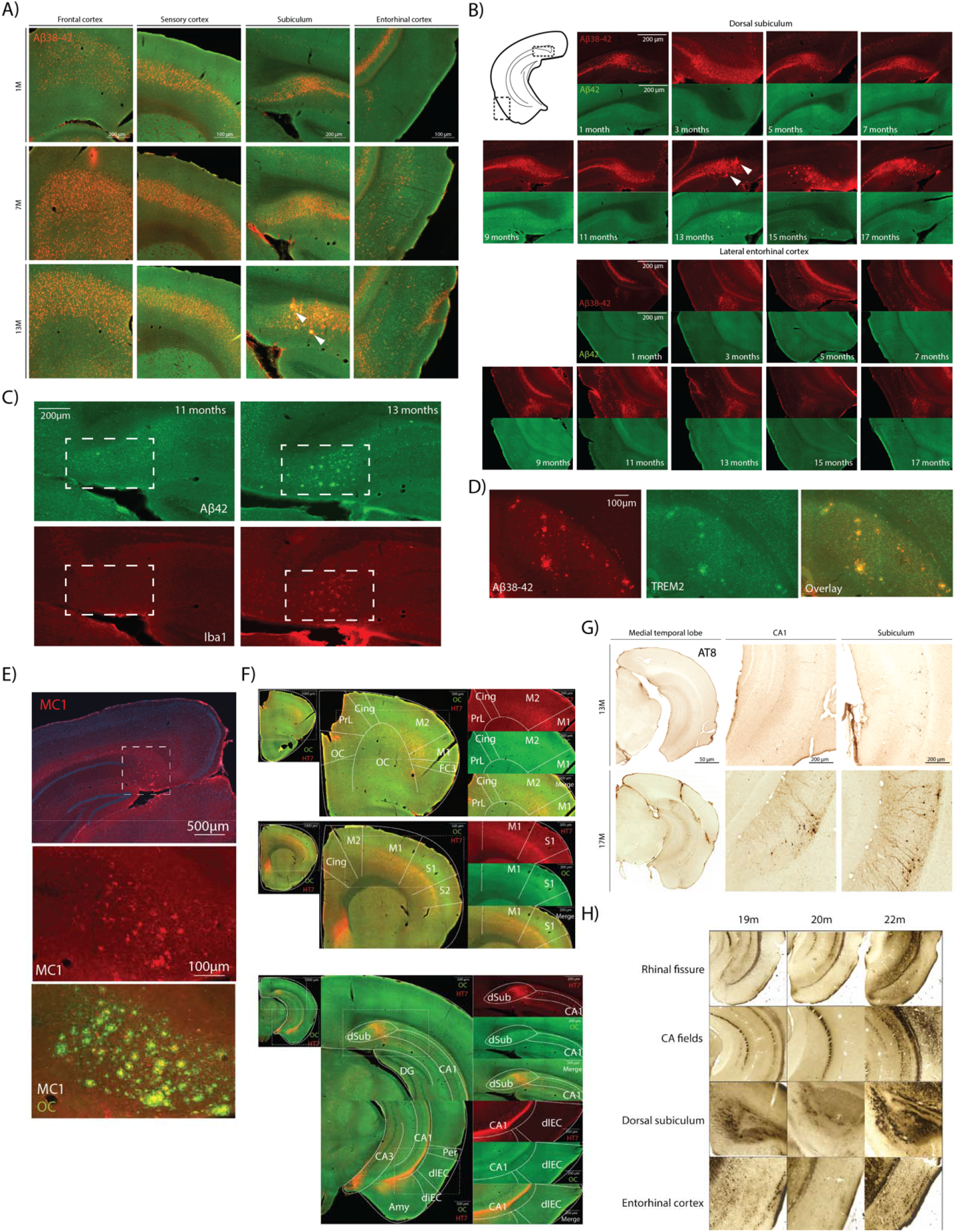
The 3xTg AD mouse model displays Alzheimer’s disease-related neuropathology. **A)** Aβ38-42 immunoreactivity in frontal cortex, sensory cortex, Sub, and EC in the 3xTg AD mouse model at 1, 7 and 13-months-of-age. Aβ38-42 (McSA1; red) and Aβ42 (IBL Aβ42; green) immunoreactivity in the 3xTg AD mouse model. Amyloid plaques immunoreactive to Aβ38-42 (McSA1 antibody) are first apparent at 13 months of age in subiculum. **B)** Aβ38-42 immunoreactivity in the 3xTg AD mouse model at various ages in the dSub and LEC. Aβ38-40 (McSA1; red) and Aβ42 (IBL Aβ42; green) immunoreactivity in the 3xTg AD mouse model. According to the ABC scoring system, scoring of diffuse amyloid plaques is assessed in the cerebral cortex, hippocampus, striatum, midbrain, brainstem, and cerebellum according to protocols established by Thal (*33*) resulting in a Thal phase 0-5, which is translated into the NIA-AA score of A0-A3. Amyloid plaques immunoreactive to Aβ38-42 (McSA1 antibody) are first apparent at 13 months of age in dSub, whereas plaques immunoreactive to Aβ42 (IBL Aβ42 antibody) are first apparent at 15 months of age in dSub. **C)** Aβ42 (IBL Aβ42; green) and Iba1 (microglial marker; red) immunoreactivity in dSub at 11 and 13-months-of-age in the 3xTg AD mouse model. **D)** Aβ38-42 (McSA1; red) and TREM2 (microglial receptor; green) immunoreactivity in dSub at 13-months-of-age. **E)** Conformation-specific tau (MC1; red) and fibrillar Aβ (OC; green) immunoreactivity in dSub at 13-months-of-age. **F)** Endogenous MAPT (HT7; red) and fibrillar Aβ (OC; green) immunoreactivity in the entire brain at 1-month-of-age in the 3xTg AD mouse model. **G)** AT8 (detects phosphorylated tau proteins at serine 202 and threonine 205 residues; DAB) immunoreactivity in the 3xTg AD mouse at 13 and 17-months-of-age. An ABC score for NFTs is determined in the trans-entorhinal area, corpora ammonis, fronto-parietal cortex, and primary visual cortex to generate a Braak stage (*34*), which is translated into the NIA-AA score of B0-B3. Most existing mouse models do not generate NFTs, so the National Institutes of Health (NIH) have developed a modified B score for p-tau pathology, including distribution of cytoplasmic neuronal tau such as pre-tangles and threads. **H)** Gallyas silver staining in the 3xTg AD mouse at 19, 20 and 22-months-of age. Abbreviations; M: month(s); Aβ: amyloid-β; Sub: subiculum; dSub: dorsal subiculum; LEC: lateral entorhinal cortex; S1: primary somatosensory cortex; S2: secondary somatosensory cortex; Olf: olfactory area; OC: orbital cortex; PrL: prelimbic cortex; Cing: cingulate cortex; M1: primary motor cortex; M2: secondary motor cortex; FC3: frontal cortexarea 3; Ins: insular cortex; CA1: cornu ammonis field 1; CA2: cornu ammonis field 2; CA3: cornu ammonis field 3; Amy: amygdala; diEC: dorsal intermediate entorhinal cortex; dlEC: dorsolateral entorhinal cortex; dSub: dorsal subiculum; PER: perirhinal cortex.

In the current study, intracellular tau immunoreactive to HT7 (endogenous microtubule-associated protein tau [MAPT]) is present in frontal and sensory areas, and the HPC as early as 1 month of age (**Fig. 3F**). The HT7 immunoreactivity appears to be less pronounced in 13-month-old mice compared to 1 month old mice, suggesting that tau levels decrease in later stages of AD. Regarding pathological tau (AT8 antibody) immunoreactivity in 13-month-old 3xTg AD mice, there is immunolabelling in CA1 and caudal subiculum at this age, but not immunolabelling in dSub (**Fig. 3G**). At 13 months of age, caudal subiculum appears to have strongest immunolabelling, and is therefore the first region involved in tau pathology in this model, at least in terms of hyperphosphorylated tau. There are NFTs present in pyramidal CA1 neurons at 18 months of age, and the brain region with the most prominent AT8 immunoreactivity is the caudal subiculum at 17 months of age. Gallyas-Braak staining demonstrates the accumulation of phosphorylated tau, such as neurofibrillary tangles and glial inclusions. Gallyas silver staining (**Fig. 3H**) in aged 3xTg AD mice (*n* = 6) between 19-and 22-months-of-age reveals NFTs throughout the entire brain parenchyma, with strongly stained inclusions in dSub and EC. However, since tau deposition does not begin in the superficial layers of LEC in this mouse model as it does in patients, a group of mice was injected with MAPT_*P301L*_ human tau (huTau) into this region to better mimic spatiotemporal spread of tauopathy.

### Fasudil and Lonafarnib infusions reduced amyloid-β and tau

As Lonafarnib has previously been shown to increase the activation of lysosomes (*26*), we hypothesized that Lonafarnib infusions would increase lysosomal associated membrane protein 1 (LAMP1) immunoreactivity after infusions in 3xTg AD mice. LAMP1 immunoreactivity was more prominent in the dSub of Lonafarnib infused mice (*n* = 2) compared to saline infused mice (*n* = 2), particularly in amyloid plaques in the dSub (**Fig. 4A**). Fasudil infusions reduced the number of intracellular Aβ-positive neurons in dSub in young 3xTg mice (**Fig. 4B**; *n* = 3; *t*_*18*_ = 2.634, *P* = 0.0169, unpaired two-tailed t-test). Lonafarnib did not affect the number of intracellular Aβ-positive neurons in dSub in young 3xTg AD mice (**Supplementary Fig. 2**). However, Lonafarnib infusions appear to have slightly reduced the number of human tau (huTau) neurons in lateral EC after injection of P301L mutated tau (**Fig. 4C**; *n* = 4; *n*.*s*.). In older 3xTg AD mice, Fasudil infusions reduced the number (*n*.*s*.) and size (*t*_*26*_ = 4.685, *P* < 0.0001, unpaired two-tailed t-test) of amyloid plaques, whereas Lonafarnib infusions significantly reduced the number (*t*_*14*_ = 3.861, *P* = 0.0017, unpaired two-tailed t-test), but not size (*n*.*s*.) of amyloid plaques in 3xTg AD mice (**Fig. 4D**; *n* = 5). In older animals, Lonafarnib infusions increased Aβ40 and t-tau levels in CSF (**Supplementary Fig. 3**). Lonafarnib infusions appear to lower MCI-positive neurons in dSub after amyloid plaque deposition in 3xTg AD mice (**Fig. 4E**; *n* = 2). Moreover, Fasudil infusions decreased the number and size of dense-core amyloid plaques, whereas Lonafarnib decreased the number but increases the size of these forms of amyloid plaques (**Supplementary Fig. 4**). Fasudil and lonafarnib infusions both decreased Aβ40 and t-tau levels in CSF (**Fig. 4F, G**).

**Figure 4.**
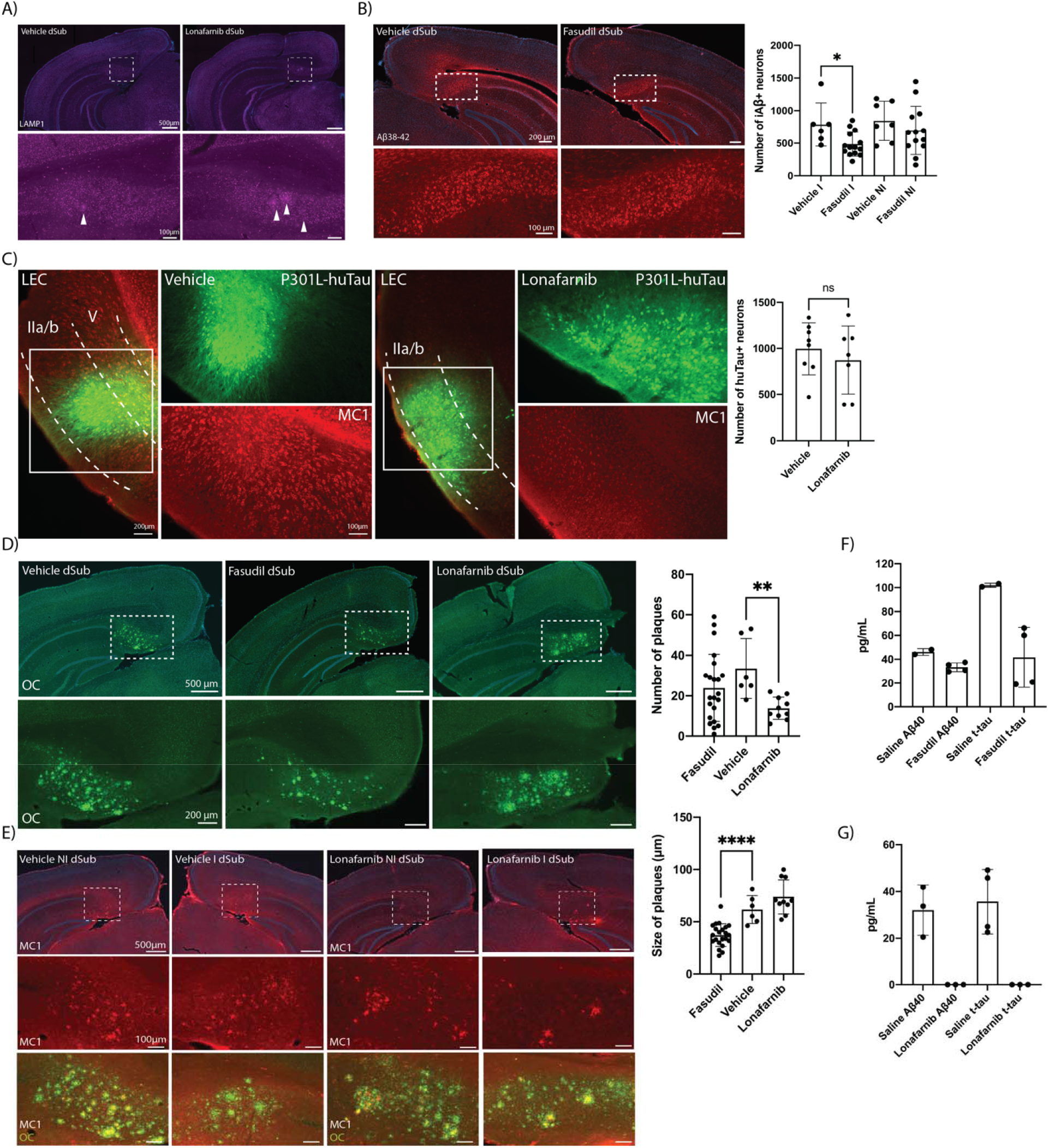
Fasudil and Lonafarnib infusions reduced Aβ and tau. **A)** LAMP1 (lysosomal marker; purple) immunoreactivity (arrowheads) in dSub in 3xTg AD mice (*n* = 4) receiving infusions of a vehicle or Lonafarnib. **B)** Aβ38-42 (McSA1; red) immunoreactivity in dSub in 3xTg AD mice (*n* = 3) receiving infusions of a vehicle or Fasudil. Mean number of intracellular Aβ+ neurons and SD displayed in bar graph, unpaired two-tailed t-test. **C)** P301L tau injected (GFP; green) 3xTg mice AD (*n* = 4) receiving infusions of a vehicle or Lonafarnib and MC1 (conformation-specific tau; red) immunoreactivity in lateral EC. Mean number of huTau+ neurons and SD displayed in bar graph, unpaired two-tailed t-test. **D)** Amyloid plaques (OC; green) in dSub of 3xTg AD mice (*n* = 5) receiving infusions of a vehicle, Fasudil or Lonafarnib. Mean number and size of amyloid plaques and SD displayed in bar graph, unpaired two-tailed t-test. **E)** MC1 (conformation-specific tau; red) and OC (amyloid plaques; green) immunoreactivity in 3xTg AD mice (*n* = 2) receiving infusions of a vehicle or Lonafarnib in infused (I) and non-infused (NI) hemispheres. **F)** Treatment with sterile saline or Fasudil on day 8 in 7-month-old 3xTg AD mice (*n* = 4). Bar graph displays mean pg/ml levels of Aβ40 and t-tau in CSF, error bars = SD. **G)** Treatment with sterile saline or Lonafarnib on day 9 in 6-month-old 3xTg AD mice (*n* = 4). Bar graph displays mean pg/ml levels of Aβ40 and t-tau in CSF, error bars = SD. Abbreviations; dSub: dorsal subiculum; Aβ: amyloid-β: iAβ: intracellular amyloid-β; LEC: lateral entorhinal cortex; huTau: human tau; NI: non-infused; I: infused.

### Co-infusion of Fasudil and Lonafarnib is more effective in reducing neuropathology

Since both drugs appeared to either reduced Aβ (Fasudil) or tau (Lonafarnib) during our pilot experiments, we did co-infusions of both drugs and assessed intracellular Aβ accumulation in the dSub of 3xTg AD mice. Co-infusions of both drugs greatly reduced the number of intracellular Aβ-positive neurons in dSub (**Fig. 5A**; *n* = 11; *U* = 3.54.5, *P* < 0.0001, Mann-Whitney U test). Infusions by *in vivo* microdialysis were more effective in reducing intracellular Aβ accumulation in the dSub compared to oral administration of the drugs (**Fig. 5B**; *n* = 3; *t*_*65*_ = 2.537, *P* = 0.0136, unpaired two-tailed t-test). Co-infusion of both drugs reduced Aβ40 and t-tau levels in CSF (**Fig. 5C**). Co-infusion of both drugs appeared to slightly improve memory when tested in a contextual memory paradigm (**Fig. 5D**; *n* = 3). Animals infused with a vehicle and those infused with drugs were able to complete morphed contextual experimental testing and were therefore tested in the Squircle context (*35*). Mice infused with drugs were slightly better at digging in the reward locations associated with training in a square and a circle compared to animals infused with a vehicle.

**Figure 5.**
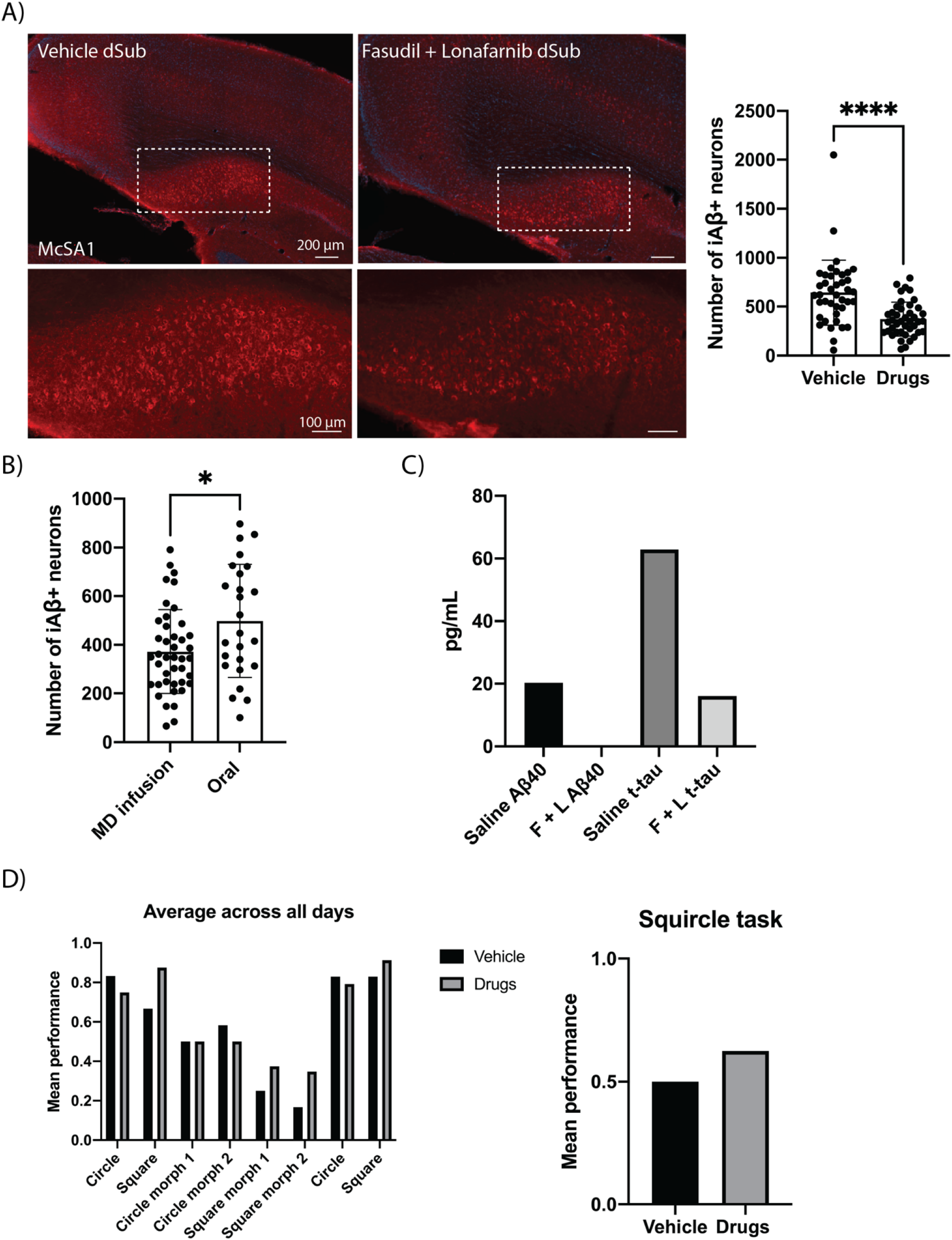
Co-infusion of Fasudil and Lonafarnib is more effective in reducing neuropathology. **A)** Aβ38-42 (McSA1; red) immunoreactivity in dSub in 3xTg AD mice (*n* = 11) receiving infusions of a vehicle, or Lonafarnib and Fasudil. Median number of intracellular Aβ+ neurons and IQR displayed in bar graph, Mann-Whitney U test. **B)** Mean number of intracellular Aβ+ neurons in dSub after infusions of Lonafarnib and Fasudil via in vivo microdialysis (*n* = 11) or oral administration (*n* = 3). Mean number of intracellular Aβ+ neurons and SD displayed in bar graph, unpaired two-tailed t-test. **C)** Treatment with sterile saline or Fasudil + Lonafarnib on day 7 in 1-month-old 3xTg AD mice (*n* = 4). Bar graph displays mean pg/ml levels of Aβ40 and t-tau in CSF. **D)** Mean performance in morphed contexts across all days and performance in the Squircle context in 3xTg mice AD (*n* =3) receiving either a vehicle or co-infusion of Fasudil and Lonafarnib. Abbreviations; dSub: dorsal subiculum; iAβ: intracellular amyloid-β; MD: microdialysis.

**Figure 6.**
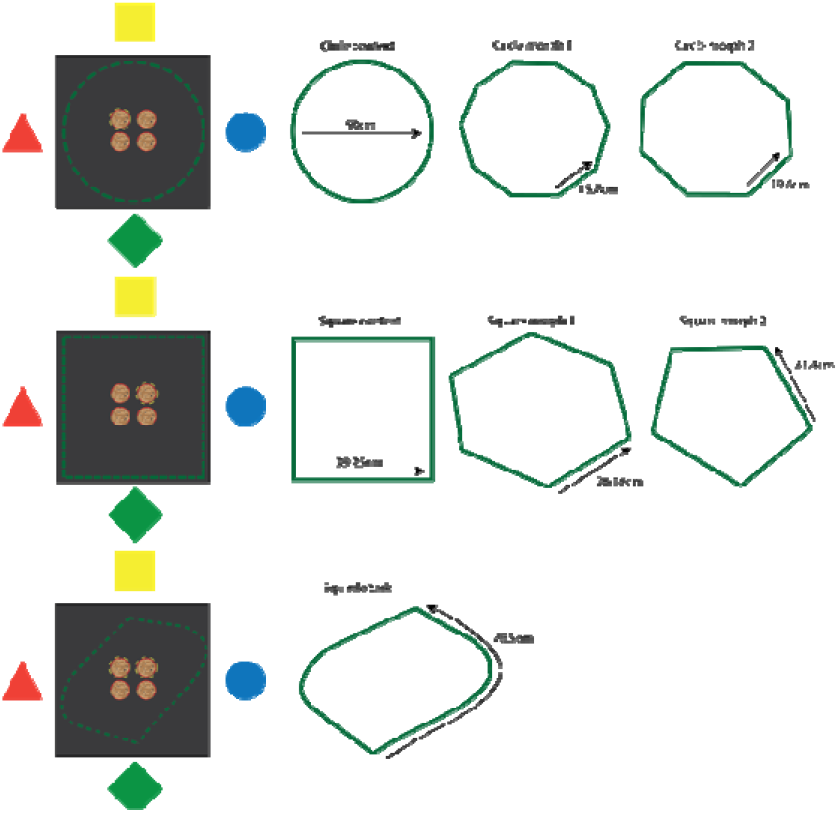
The Squircle task. Mice were initially taught to associate a specific reward location in a square and a circle context. The reward was embedded in one of four cups with ginger-scented bedding. A trial was assessed as correct if the mice dug in the reward location associated with each chamber. If they passed the training phase (66.6% correct digging), their contextual memory performance was tested in morph chambers: decagon (circle morph 1), octagon (circle morph 2), hexagon (square morph 1), an a pentagon (square morph 2). Mice were tested in morph chambers for 4 days, with 8 sessions a day. If they were able to complete the morph testing, they were tested in a Squircle chamber on the fifth day. Since the Squircle equally resembles a square and a circle context, a trial was considered correct if the mouse dug either of the reward locations attributable to the square-or circle context. If so, it was attributed as contextual association, if not, it was attributed as visual learning.

## DISCUSSION

Before proceeding with drug administration, it was important to establish how well the specific mouse model recapitulates key hallmarks of human AD pathology. Our findings support that the 3xTg AD mouse model captures AD-related neuropathology, with intracellular Aβ appearing first in the HPC, where subsequent amyloid plaques are deposited at later stages. According to previous characterizations, intracellular Aβ immunoreactivity should be present in CA1, amygdala and neocortex at 4 months of age (*22, 36*), whereas here we found it to be observed in frontal and sensory areas, the HPC, and parts of the cerebellum as early as 1 month of age. Earlier characterizations of this mouse line have observed amyloid plaque deposits in the frontal cortex at 6 months of age (*22, 36*). The present findings are contradictory to earlier characterization, as amyloid plaques are observed first at 13 months of age in the dSub. Amyloid plaques deposited in dSub of the 3xTg AD mouse model are associated with TREM2-related immunoreactivity. Moreover, MC1 immunoreactivity within OC-positive amyloid plaques gives support to the interplay between Aβ and tau in AD pathophysiology (*37*). Previous findings have shown that aggregates of hyperphosphorylated tau in this mouse line is present in CA1 at 12-15 months of age (*22*), whereas here we found it to be present in frontal and sensory areas, and the HPC as early as 1 month of age. This mouse model harbors endogenous tau that leads to subsequent NFT development in subiculum in older ages. There are NFTs present in pyramidal CA1 neurons at 18 months of age, however there are fewer NFTs than reported by Oddo and colleagues (*22*). Overall, this mouse model presents as a valid model for studying the synergistic effects of Aβ and tau in AD.

Treatment with Fasudil reduced intracellular Aβ in the brain and CSF Aβ, as well as the number and size of amyloid plaques in HPC. This is in line with the notion that when the Wnt signaling pathway is inhibited it reduces Aβ production (*38-40*). Treatment with this drug also reduced t-tau levels in CSF. Lonafarnib, on the other hand, reduced conformation-specific tau in LEC following injections of P301L huTau and the number of amyloid plaques in HPC. This lends support to findings that suggest that the expression of autophagic proteins decline with age, which may impact neurodegeneration in AD (*41, 42*). Treatment with this drug also reduced Aβ and t-tau levels in CSF. On the other hand, we can only speculate that the finding of larger amyloid plaques after treatment with Lonafarnib could be due to the increase of intracellular digestion, resulting in a larger expansion of amyloid plaques (*43*). In older animals, treatment with Lonafarnib increased Aβ levels in CSF. Both drugs targeted dense-core amyloid plaques, and these depositions are associated with microglial activation, subsequent neurodegeneration, and cognitive decline in human patients (*44, 45*). Intriguingly, we found that co-infusion of both drugs greatly reduced intracellular Aβ in HPC and positively affected contextual memory performance in mice.

Early postmortem studies of brains from AD patients suggested that initial accumulation of Aβ occurs in the neocortex with subsequent spread of aggregates to deeper structures such as those in the medial temporal lobe (*33*). However, during postmortem studies, reports are unable to document Aβ deposition over time prior to the development of AD-related dementia. Recent advances in positron emission tomography (PET) imaging have enabled longitudinal human Aβ-imaging studies that confirm the importance of cortical Aβ in the diagnosis and prediction of progression of cognitive impairment (*46, 47*), although they have largely not examined deeper structures for Aβ load and spread (*47, 48*). Therefore, a systematic and comprehensive approach to studying Aβ deposition may reveal unexplored Aβ-related changes in the brain of AD patients (*49*). In line with this, recent findings suggest that subcortical memory network hubs may be critically susceptible to pathological changes that occur in AD and that changes within these networks may contribute to cognitive decline (*50*). Regarding Aβ spread, the dSub, and other hippocampal regions, are involved prior to the EC in the 3xTg AD mouse model. Our characterization results also suggest that early intracellular Aβ might be in monomeric or fibrillar forms, and not oligomeric, and the latter forms are better correlated with disease severity than are amyloid plaques containing insoluble Aβ fibrillary species (*51, 52*). These latter results are in line with the notion that Aβ accumulates over decades in human patients without affecting cognitive decline prior to the deposition of amyloid plaques (*53*).

Several factors suggest that simultaneous treatment with Fasudil and Lonafarnib would be more beneficial than either drug alone. For instance, amyloidogenic processing of APP occurs in endosomes, following clathrin-mediated endocytosis (*54*), and endosomal trafficking of familial AD forms of *APP* has been shown to differ from that of wild-type *APP* (*55*). Interestingly, Wnt-PCP signaling is also tightly linked to clathrin-mediated endocytosis (*56*). This follows our findings of reduced dense-core amyloid plaques in mice after Fasudil treatment. Moreover, researchers have found that clathrin-positive endosomes containing Aβ are involved in a process of LC3-associated endocytosis (*57*). This endocytic pathway was found to be a critical regulator of microglia for removal of protein aggregates in mice modelling AD. Mice deficient in LC3-associated endocytosis displayed accelerated neurodegeneration, impaired neuronal signalling, and memory deficits. This seems to be in line with our characterization of the 3xTg AD mouse model and our observation of TREM2-positive microglia clustering around amyloid plaques.

Push-pull microdialysis offers a powerful method for the simultaneous collection of CSF and infusion of disease-targeting drugs, however, there are potential limitations that should be considered. Recovery rates, which are inversely proportional to the perfusate flow rate (*58, 59*), for all four CSF biomarkers (Aβ40, Aβ42, t-tau, p-tau) were relatively low compared to previous findings (*60*) for longitudinal CSF biomarkers in the 3xTg AD mouse model, which was likely due to a relatively fast sampling rate (1 μL/minute). We have previously found that a sampling rate of 0.2 μL/minute was ideal for the recovery of CSF Aβ and tau (*60*). However, due to the repeated infusions of drugs in animals in the current study, a faster sampling rate was used to reduce infusion time per animal as well as to administer the drugs as one dose, rather than as a slow diffusion process over many hours. Additionally, as CSF t-tau levels generally increased in the first few days of sampling, it is likely that the removal and insertion of the microdialysis dummy and sampling probe resulted in minor tissue damage at the start of sampling. This is consistent with the fact that in response to acute brain injury, CSF t-tau concentrations increase in the first few days following injury, then reduce over time (*61*). However, we have previously shown that long-term implantation of our microdialysis probe does not lead to increased neuroinflammation in 3xTg AD mice (*60*).

Moreover, intracerebral microdialysis is an invasive technique, so it is currently only used in patients requiring neurocritical care, neurosurgery, or brain biopsy (*62*). A leaky BBB often precedes amyloid plaque formation in AD patients (*63*), and this is also observed in 3xTg AD mice (*64*). Enlarged ventricles and concomitant brain atrophy could be a possible indicator of AD disease progression (*65*). In line with this, a study has shown that in the 3xTg AD mouse model, ventricles become enlarged compared to control mice already at 2 months-of-age (*66*). Moreover, in our study, and most preclinical translational AD research, there is an increased focus on familial rather than sporadic forms of AD. Whereas most AD cases are sporadic, transgenic animal models of AD are based on genetic mutations found in familial AD, specifically in the *APP*, presenilin 1 (*PSEN1*) and presenilin 2 (*PSEN2*) genes (*67*). This limitation, however, should not detract from the value of the current study and other studies using transgenic models, as the continuum of sporadic and familial forms of AD are very similar, except for age-of-onset (*67*).

### Conclusions and future directions

Due to the current failure in disease-targeting drugs to attenuate AD neuropathology, there is an urgent need for novel therapeutic approaches. Here we repurposed two FDA-approved drugs, Fasudil and Lonafarnib, with independent actions on biochemical events leading to AD pathology. Treating 3xTg AD mice with both drugs simultaneously was more effective at reducing intracellular Aβ and led to better contextual memory performance. However, an unexpected finding was that Lonafarnib treatment increased amyloid plaques and CSF Aβ in old animals, suggesting that activating the endosomal-lysosomal system may inadvertently increase Aβ pathology if administered too late in the AD continuum. In line with this, it may be interesting to infuse Lonafarnib prior to over-expression of huTau in lateral EC. Future studies should aim to longitudinally infuse both drugs and examine the effect on AD pathophysiology. Furthermore, there is currently an urgent need within the field to develop blood-based biomarkers which are inexpensive and minimally invasive, which can detect early neuropathological changes in AD (*68*). It is important to note, however, that blood biomarkers have lower sensitivity and specificity than CSF biomarkers, and this can be attributed to the fact that the BBB prevents diffusion of analytes into the blood via a filtering mechanism (*69*), and there is currently no approved blood biomarker for AD. Taken together, our findings lend support to the application of repurposed drugs to attenuate Alzheimer’s disease neuropathology at various therapeutic time windows in preclinical models and thus obtain insights into mechanisms of action that may attenuate neuropathology in AD patients.

## MATERIALS AND METHODS

### Animals

Thirty 3xTg AD mice and two control B6129 mice were included in these experiments. These mice contain three mutations associated with familial Alzheimer’s disease (*APP*_*Swe*_, *MAPT*_*P301L*_, and *PSEN1*_*M146V*_). The 3xTg AD mouse model is based on a C57BL/6;129X1/SvJ;129S1/Sv (Jackson Laboratory, Bar Harbor, ME, USA) background. The model exhibits both amyloid plaques and NFTs, and cognitive impairment appear at 4 months of age in these mice.

The donating investigators of the 3xTg AD mouse model communicated to Jackson Laboratories that male transgenic mice may not exhibit the phenotypic traits as originally described (*36*). Therefore, only female mice were included in these experiments. All housing and breeding of animals used was approved by the Norwegian Animal Research Authority and is in accordance with the Norwegian Animal Welfare Act §§ 1-28, the Norwegian Regulations of Animal Research §§ 1-26, and the European Convention for the Protection of Vertebrate Animals used for Experimental and Other Scientific Purposes (FOTS ID 21061). The animals were kept on a 12h light/dark cycle under standard laboratory conditions (19-22°C, 50-60% humidity), and had free access to food and water.

### Microdialysis guide implantation surgeries

Surgeries were based on our earlier established protocol (*60, 70*), but in brief, implantation surgery was performed to insert microdialysis guide cannulas (CMA 7; CMA Microdialysis AB, Kista, Sweden) into the lateral ventricle of mice. Mice were anesthetized with isoflurane gas (4 % induction and 1.5-3 % maintenance; IsoFlo vet., Abbott Laboratories, Chicago, IL, USA) prior to being fixed in a stereotaxic frame (Kopf Instruments; Chicago, IL, USA). Prior to making any incisions, Marcain (0.03-0.18 mg/kg; Aspen Pharma, Ballerup, Denmark) was injected subcutaneously into the scalp and Metacam (5 mg/kg; Boehringer Ingelheim Vetmedia, Copenhagen, Denmark) and Temgesic (0.05-0.1 mg/kg; Invidor UK, Slough, Great Britain) were administered subcutaneously for intraoperative pain relief. Equal heights of bregma and lambda were measured to ensure that the skull was level for each animal (with ±0.1 mm tolerance), as well as 2 points equally distant from the midline. After levelling the skull, the stereotaxic coordinates were derived to target the lateral ventricle (A/P -0.1 mm, M/L +1.2 mm, D/V -2.75 mm). The microdialysis guide cannula was attached to the stereotaxic frame using a guide clip and connection rod for the clip (CMA Microdialysis AB, Kista, Sweden). The skull was drilled through at these coordinates and the guide cannula was slowly lowered into the drilled hole. The guide cannula was attached to the skull with super glue and dental cement (Dentalon Plus; Cliniclands AB, Trelleborg, Sweden). Post-surgery, Metacam and Temgesic were administered within 24 hours. The guide cannula was implanted into the right hemisphere of all animals, as we did not observe any lateralization of pathology in the brains of 3xTg AD mice (**Supplementary Fig. 5**).

### Stereotaxic viral injections of P301L tau

Mice were treated identically to microdialysis guide implantation surgeries, up until deriving stereotaxic coordinates. A craniotomy was made at 0.5 mm anterior to lambda and ∼4 mm lateral (dependent on animal weight) to the midline. A Hamilton microsyringe (Neuros 32-gauge syringe, 5μl, Hamilton company, Nevada, USA) was lowered vertically into the brain to a depth ∼3.6 mm (dependent on animal weight) from the surface, and 2000-2500 nL of viruses was injected using a microinjector (Nanoliter 2010, World Precision Instruments Inc., USA). We used AAV8-P301L Tau-2a-GFP for injections into layer II of lateral EC (generated in house by Dr. Nair). The microsyringe was kept in place for 5 minutes prior and after the injection, to minimize potential upward leakage of the viral solution. Metacam was given within 24 hours post-surgery. Animals were implanted with microdialysis guide cannulas two months following injections.

### Push-pull microdialysis apparatus and sampling

Push-pull microdialysis was conducted as previously described (*60, 71*), but in brief a refrigerated fraction collector (CMA 470) was set to 6 °C for the storage of collected CSF in 300 μl low-retention polypropylene plastic vials (Harvard Apparatus, Cambridge, MA, USA). Fluorinated ethylene propylene (FEP) peristaltic tubing (CMA Microdialysis AB, Kista, Sweden) was placed inside each plastic vial for collection and connected to the cassette of the peristaltic roller pump (Reglo ICC Digital). This peristaltic FEP tubing was connected to the outlet side of microdialysis probes (β-irrigated 2 mDa microdialysis probe; CMA 7; CMA Microdialysis AB, Kista, Sweden) with a polyethersulfone 2 mm membrane with tubing adapters bathed in 75 % ethanol. FEP tubing (CMA Microdialysis AB, Kista, Sweden) was connected to each microsyringe. The FEP tubing was then connected to the inlet part of the microdialysis probes. Transparent cages were prepared with 1.5 cm of bedding, filled water bottles, and treats. Saline or drugs loaded inside a gastight microsyringe (CMA Microdialysis AB, Kista, Sweden), which was placed into a syringe pump (CMA 4004). The ‘dead volume’ of the FEP outlet tubing (1.2 mL/100mm) was calculated. 100 cm of FEP outlet tubing was used, and therefore the first 12 mL sampled from each animal were discarded. Prior to inserting the microdialysis probes into the guide cannula, the probe was conditioned in 75 % ethanol for better recovery of analytes. At the conclusion of microdialyte sampling, the vials of 60 μl CSF were centrifuged and kept at -80 °C until the samples were analyzed with multiplex ELISA.

### Drug infusions into the lateral ventricle

Fasudil is a promising drug candidate for AD as it has previously been given clinical approval for the treatment of cerebral vasospasm and is therefore safe to use in humans. Previously, researchers have administered Fasudil at 10mg/kg daily using DMSO as a vehicle into the lateral ventricle for 14 days (*24*). However, DMSO can damage the BBB, mitochondria and can cause apoptosis (*72*) and therefore all drugs used in experiments were diluted in sterile saline for delivery. Because we had a less effective delivery vehicle, Fasudil (Selleck Chemicals, Houston, TX, USA) in powder form (10mM) was diluted in sterile saline for a final concentration of 50mg/kg and stored at -80□ between infusion experiments.

Previous research has administered Lonafarnib, which is also approved for cancer treatment in humans, orally at 80mg/kg for five days on and five days off for a total of 10 weeks (*26*). For the drug to diffuse into the brain parenchyma, the same dosage was infused into the lateral ventricle in the current experiments. Lonafarnib (5 mM; Cayman Chemical, Ann Arbor, MI, USA) in powder form was dissolved in sterile saline at 60□ to a final concentration of 80mg/kg and stored at -20□ between infusion experiments.

All dosages in ml were calculated using; dosage (mg) / concentration (mg/ml) = dose x ml) and were infused at a volume of 60μl at a rate of 1 μl/min using saline as a control vehicle. In one group of animals (*n* = 3), 0.6 ml of 80 mg/kg Lonafarnib and 50 mg/kg Fasudil were mixed into baby porridge (Nestle) for oral delivery.

### Contextual memory testing

Starting 5 days before the experiment, animals were taught to dig in a brain cup for a food reward (Nestle chocolate loopie) in their home cage by providing them once daily with the reward gradually buried deeper under scented bedding. To motivate each mouse to learn the contextual paradigm, they were handled, and food deprived for five days prior to training and experimental testing.

The training of the task was conducted in a square (4 × 29.25 cm) and circle (157 cm circumference) chamber built out of rectangular Legos (2 × 1 cm, Lego, Billund, Denmark). The testing of the task was conducted in a hexagon (6 × 26.16 cm), octagon (8 × 19.63 cm), and Squircle (2 × 78.5 cm) chambers built out of rectangular Legos (2 × 1 cm). All chambers were 15 cm tall. The same base was used for all chambers, and plastic cups were embedded in the base. The plastic cups contained odor-masked bedding, consisting of 1 g of odor mask (ground ginger) for every 100 g of bedding. Testing in all chambers occurred in the same location in the experimental room. The chambers were surrounded by a square black curtain with rounded corners, and were uniformly lit from overhead. Four distal cues (a yellow square, a red triangle, a blue circle, and a green diamond all in the same size proportion) were placed on all 4 sides of the curtains. All chambers were built in proportion to each other. All rewards were mirrored in the square and circle chamber, i.e., if the reward location was the top right corner in the square, it would be the top left corner in the circle.

A pilot experiment showed that mice could discriminate the square and circle reward locations to a performance criterion after 16 training trials. Thus, all experiments began with a training phase consisting of four training trials per chamber per day for 2 days, with successive trials alternated across chambers (8 trials total in the square chamber and 8 trials total in the circle chamber). During this training, mice were taught to search for a reward (Chocolate Cheerios Cereal, Nestle, Vevey, Switzerland), which was visible for the first two training trials per chamber and buried in the remaining training trials. The correct reward location was assessed by digging, and after correctly digging 66.67% of the time, animals were ready for testing. If animals dug in the wrong location, the trial ended. If the animal did not dig after 5 minutes, the trial ended. In the square chamber, the reward was always located in the right-hand corner, whereas in the circle chamber, the reward was always located in the left-hand corner. These locations were counterbalanced across animals; however, for all analyses and figures, the percentage of searches at each corner are reflected such that the correct corner is the same for all animals.

All experiments began with a training phase consisting of four training trials per chamber per day for 2 days, with successive trials alternated across chambers (8 trials total in the square chamber and 8 trials total in the circle chamber). Animals were disoriented before the start of every trial. To disorient an animal, it was placed inside the hands of the experimenter. The experimenter slowly rotated their hands roughly four full clockwise then four full counterclockwise revolutions. The animal was then carried in the experimenter’s hands to the chamber. To ensure that the animal used distal cues, it was placed in the chamber in proportion to each side of the chamber with a 90° clockwise rotation before each trial, counterbalanced so that all orientations relative to the room were experienced equally often. The chambers were cleaned with ethanol at the end of each trial to remove odor trails. The intertrial interval was 5 min.

Following training, animals were tested in eight sessions per day for 4 days. The first two trials consisted of animals being tested in the square and circle chamber with rewards. In trials 3 to 6, the animals were tested in two morphed chambers (a morphed square and circle chamber). The order of the morphed chambers was randomized across days and balanced across animals. In trials 7 and 8, the animals were again tested in the square and circle chambers with rewards in order to not lose motivation. If an animal was able to perform the tasks up until day 4, they were tested in the Squircle context on the fifth day. During reward trials, mice were removed from the apparatus after they had found the reward. During unrewarded trials, they were removed after their first dig, or after 5 min (whichever came later). Digs were counted whenever an animal removed bedding from a cup by using one or both paws.

Dig locations and time spent in these locations were calculated using ANY-maze video tracking system (Stoelting Europe) via an overhead, centrally located camera (DMK 22AUC03 USB 2.0 monochrome industrial camera, The imaging Source Europe, Germany). Paired sample t-test were used to assess the number of correct digs, and the difference between cohorts. All reported statistics are based on two-tailed significance tests. Animals had access to water ad libitum, but to increase motivation to participate in the task, they were maintained at 90-95% of their free-feed weight.

### Proteomic analysis of amyloid-β and tau concentrations in CSF

The MILLIPLEX® MAP human amyloid beta and tau magnetic bead panel 4-plex ELISA kit (Millipore, Burlington, MA, USA) and the Bio-Plex 200 System instrument (Biorad, Hercules, CA, USA) were used to assess simultaneously the concentrations of Aβ_40_, Aβ_42_, t-tau, and p-tau (Thr181) in CSF samples. The samples were undiluted and analysed in duplicates. The lower limit of quantification (LLOQ) is shown in **Supplementary Fig. 6**.

### Tissue processing and immunohistochemistry

Mice were administered a lethal dose of sodium pentobarbital (100 mg/ml; Apotekforeningen, Oslo, Norway) and transcardially perfused with Ringer’s solution followed by paraformaldehyde (PFA, 4 %; Sigma□Aldrich) in 125 mM phosphate buffer (PB). Brains were extracted and fixed for a minimum of 24 hours in PFA at 4 °C and transferred to a 2 % DMSO solution prepared in PB for 24 hours at 4 °C. Brains were sectioned coronally at 40 μm on a freezing microtome (Microm HM430, ThermoFisher Scientific, Waltham, MA, USA). An incision was made in the non-implanted hemisphere for visualization of the control hemisphere. Immunohistochemical processing was conducted on tissue, see (*73*) for detailed protocol, and **Supplementary Table 1** for antibodies used.

Previous research has indicated that a differential microtubule-associated protein 2 (MAP2) immunolabeling pattern can distinguish dense-core amyloid plaques from diffuse amyloid plaques using DAB (Sigma-Aldrich, St. Louis, MO, USA) as a chromogen (*74*).

One series of each brain were dehydrated in ethanol, cleared in xylene (Merck Chemicals, Darmstadt, Germany) and rehydrated before staining with Cresyl violet (Nissl; 1g/l) for 3 minutes to verify probe placement. The sections were then alternatively dipped in ethanol–acetic acid (5 ml acetic acid in 1l 70 % ethanol) and rinsed with cold water until the desired differentiation was obtained, then dehydrated, cleared in xylene and coverslipped with entellan containing xylene (Merck Chemicals).

Sections were scanned using a Mirax-midi scanner (objective 20X, NA 0.8; Carl Zeiss Microscopy, Oberkochen, Germany), using either reflected fluorescence (for sections stained with a fluorophore) or transmitted white light (for sections stained with Cresyl Violet; Nissl) as the light source.

### Quantification of amyloid-β in hippocampus and tau in entorhinal cortex

Series of sections were chosen randomly and coded to ensure blinding to the investigators. The number of cells containing intracellular Aβ, tau, and amyloid plaques, in dSub and lateral EC of infused and non-infused hemispheres of vehicle-infused 3xTg AD mice (*n* = 10) and drug-infused mice (*n* = 20) was estimated with Ilastik (*75*) using the Density cell counting workflow. dSub and lateral EC was delineated using cytoarchitectonic features in sections with Nissl (Cresyl violet) stain, based on The Paxinos & Franklin mouse atlas (*76*). The same surface area and rostrocaudal levels of each brain region was selected, and at least 6 slices were used for each hemisphere. Damaged regions of brain slices were cropped out prior to quantification as this could result in false-positive antibody expression.

### Statistics

Effect size (Cohen’s D) was calculated based on initial experiments between an experimental group infused with Fasudil and a control group infused with a vehicle and the resulting effects on intracellular Aβ-positive neurons in dSub. Based on each group consisting of *n* = 4 animals, a large effect size of 1.57 was calculated (*77*). Most of the dataset we normalized (Shapiro-Wilk test) and therefore two-tailed, unpaired t-tests were used to compare mean differences. For the minor parts of the dataset that were non-normalized, nonparametric statistical tests were used (Mann-Whitney U). Correlations between drug- and vehicle-infused CSF protein levels were tested with the Pearson product moment correlation. All statistical tests and graphs were made in Prism 9 (GraphPad Software Inc., CA, USA).

## Acknowledgments

Thank you to Nora Cecilie Ebbesen for allowing us to reproduce the illustration of the Squircle task.

## Author contributions

Conceptualization: CB, IS

Methodology: CB, JBJ, AS, IS

Investigation: CB, MH

Visualization: CB, MH

Funding acquisition: CB, AS, IS

Project administration: CB, AS, IS

Supervision: AS, IS

Writing – original draft: CB

Writing – review and editing: CB, MH, JBJ, AS, IS

## Data and materials availability

All data are available in the main text or the supplementary materials.

## SUPPLEMENTARY MATERIALS

**Supplementary Figure 1.**
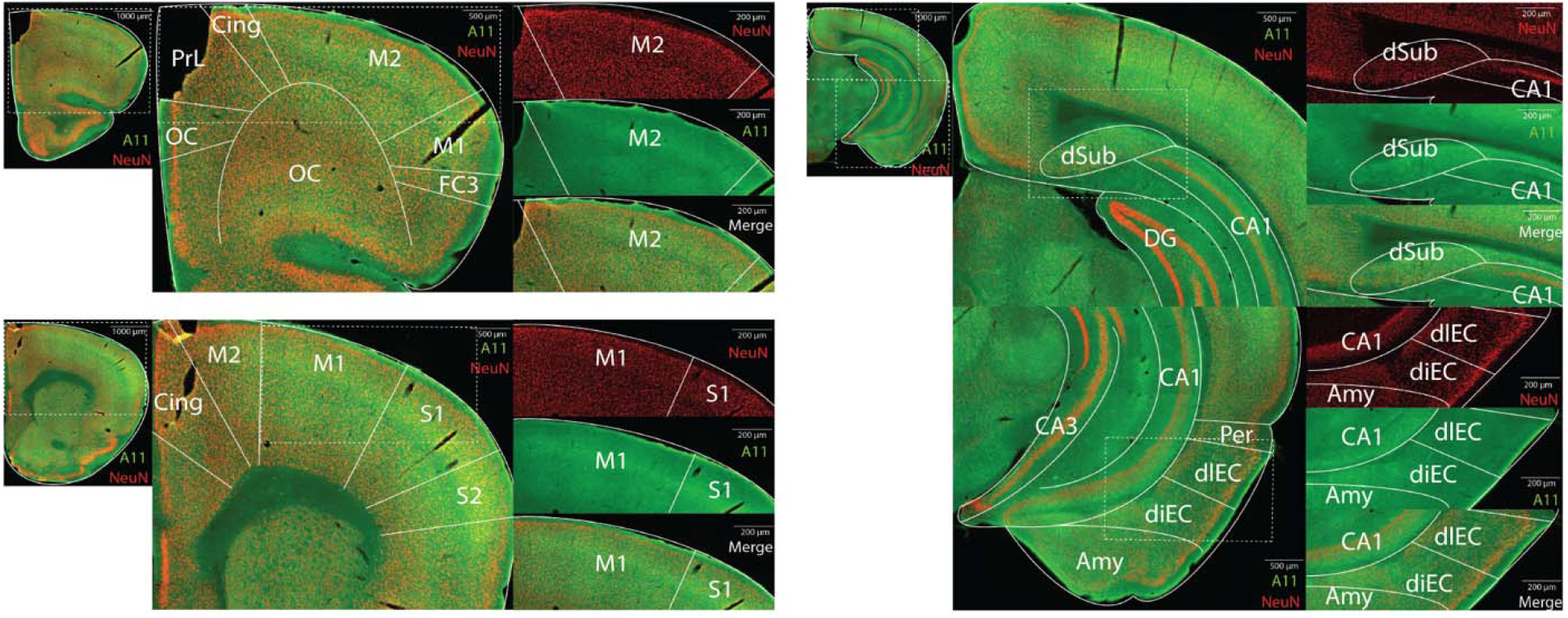
Oligomeric Aβ at 1-month-of-age in the 3xTg AD mouse model. A11 (oligomeric Aβ specific; green) and NeuN (nuclei specific; red) immunoreactivity in the 3xTg AD mouse model at 1-month-of-age. At this age, there is little-to-none A11 immunoreactivity in frontal and sensory areas of the brain (left), as well as in the hippocampal and parahippocampal region (right). Abbreviations; S1: primary somatosensory cortex; S2: secondary somatosensory cortex; Olf: olfactory area; OC: orbital cortex; PrL: prelimbic cortex; Cing: cingulate cortex; M1: primary motor cortex; M2: secondary motor cortex; FC3: frontal cortexarea 3; Ins: insular cortex; CA1: cornu ammonis field 1; CA2: cornu ammonisfield 2; CA3: cornu ammonis field 3; Amy: amygdala; diEC: dorsal intermediate entorhinal cortex; dlEC: dorsolateral entorhinal cortex; dSub: dorsal subiculum; PER: perirhinal cortex.

**Supplementary Figure 2.**
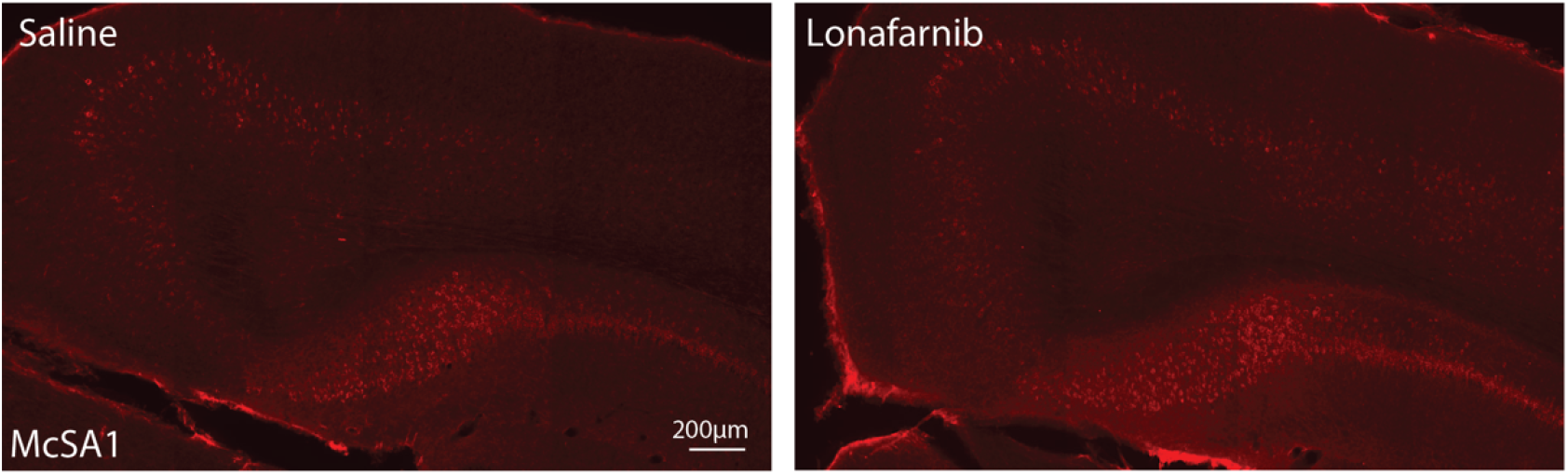
Intracellular Aβ in dSub after Lonafarnib infusions.

**Supplementary Figure 3.**
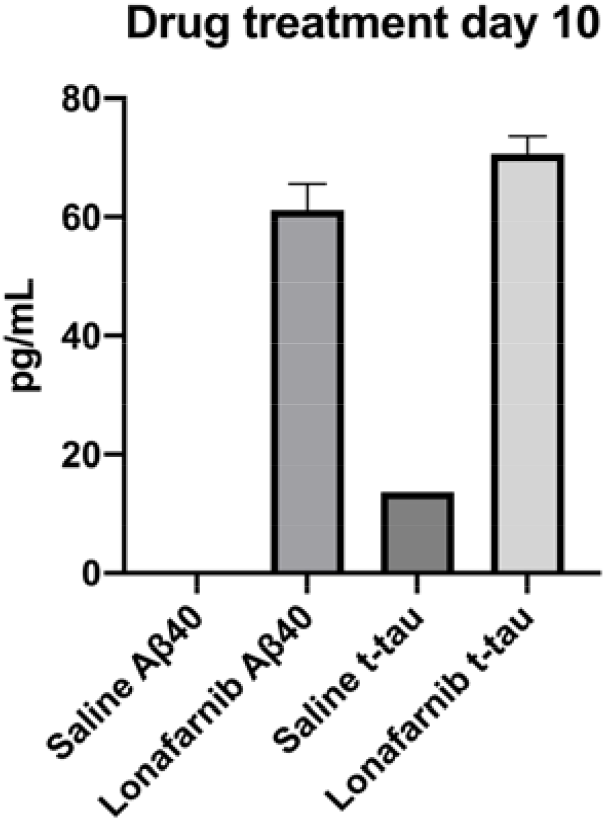
CSF Aβ40 and t-tau levels after Lonafarnib infusions in old mice. Drug treatment day 10, N = 4.

**Supplementary Figure 4.**
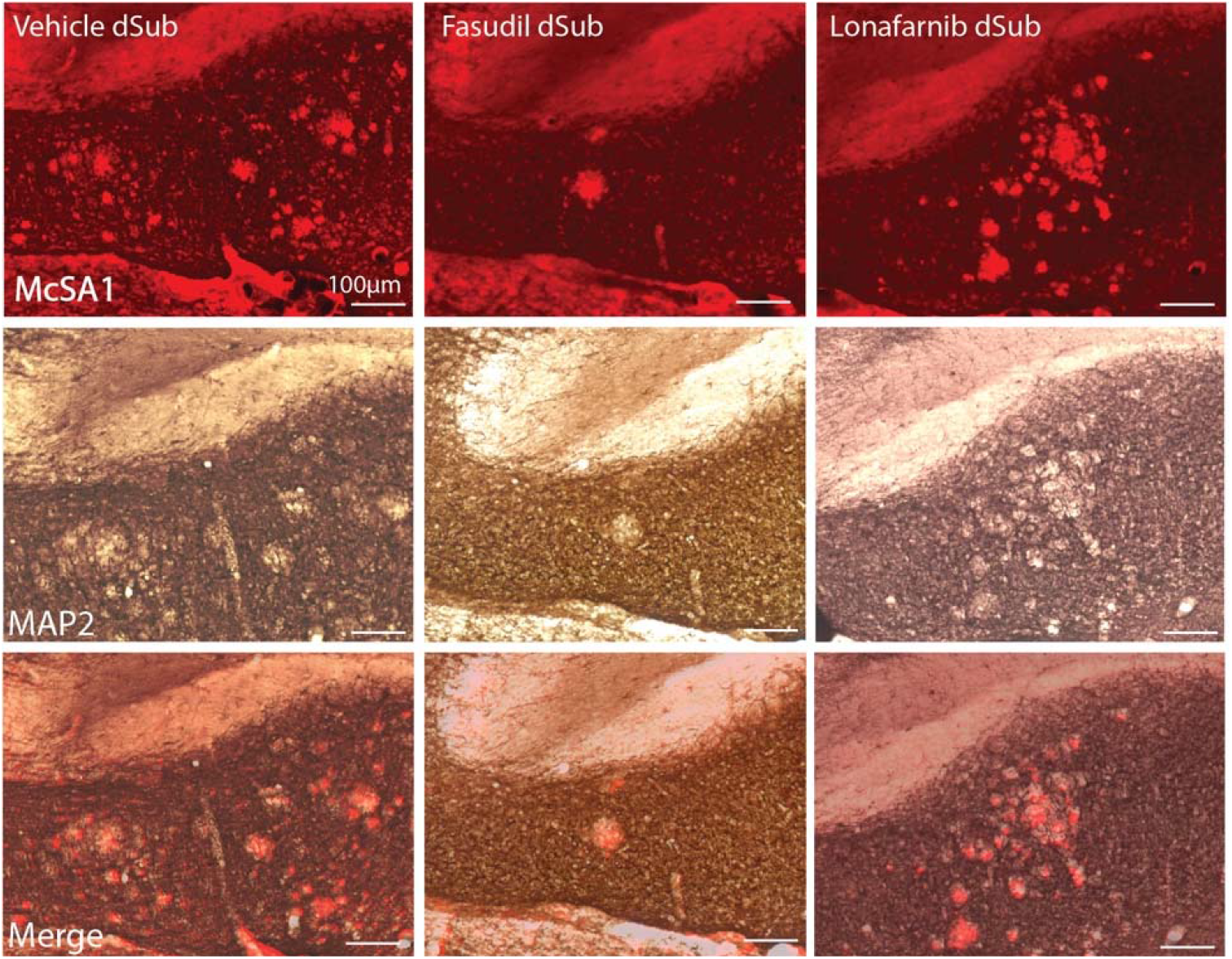
Dense-core amyloid plaques after Fasudil and Lonafarnib infusions.

**Supplementary Figure 5.**
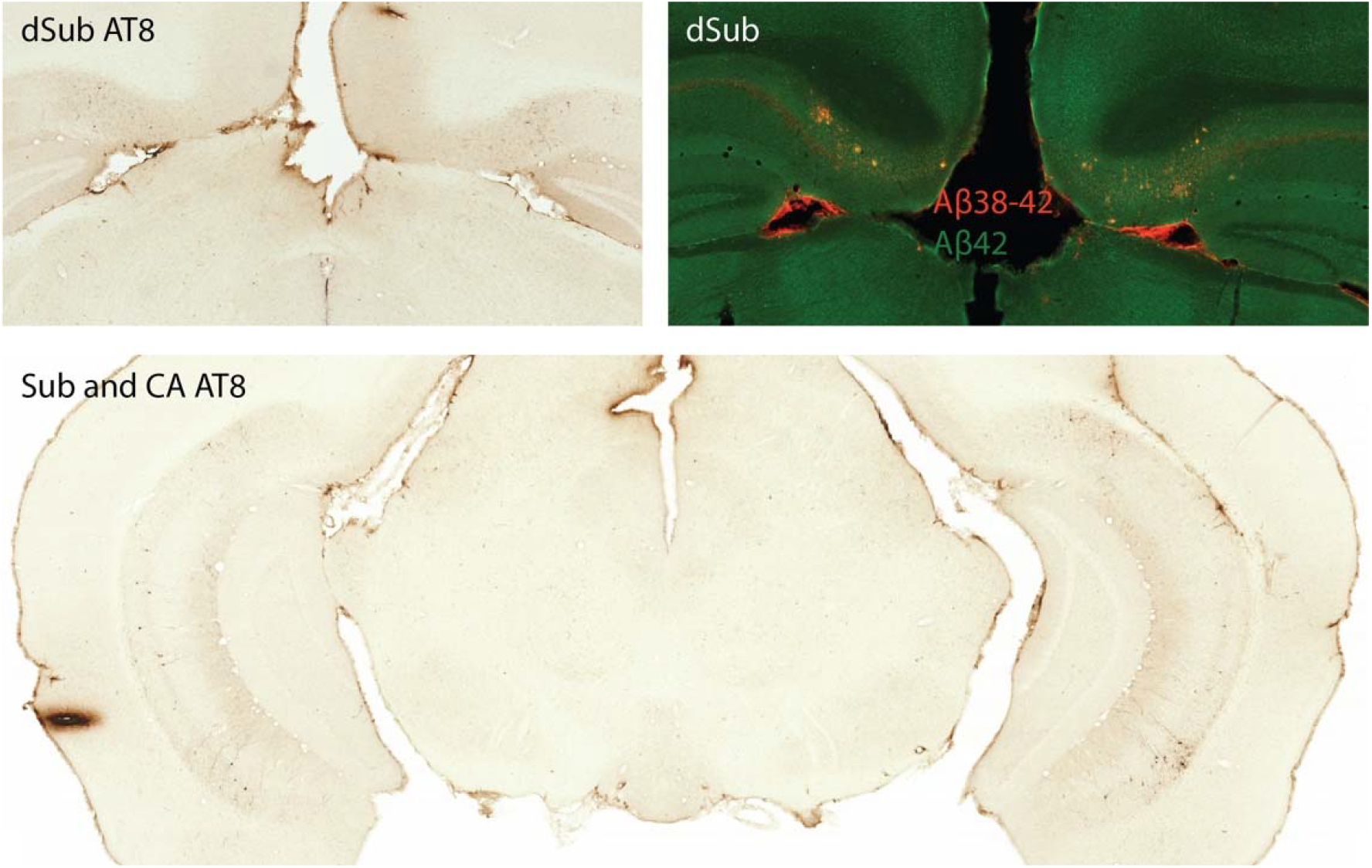
No lateralization of pathology in 3xTg AD mice.

**Supplementary Figure 6.**
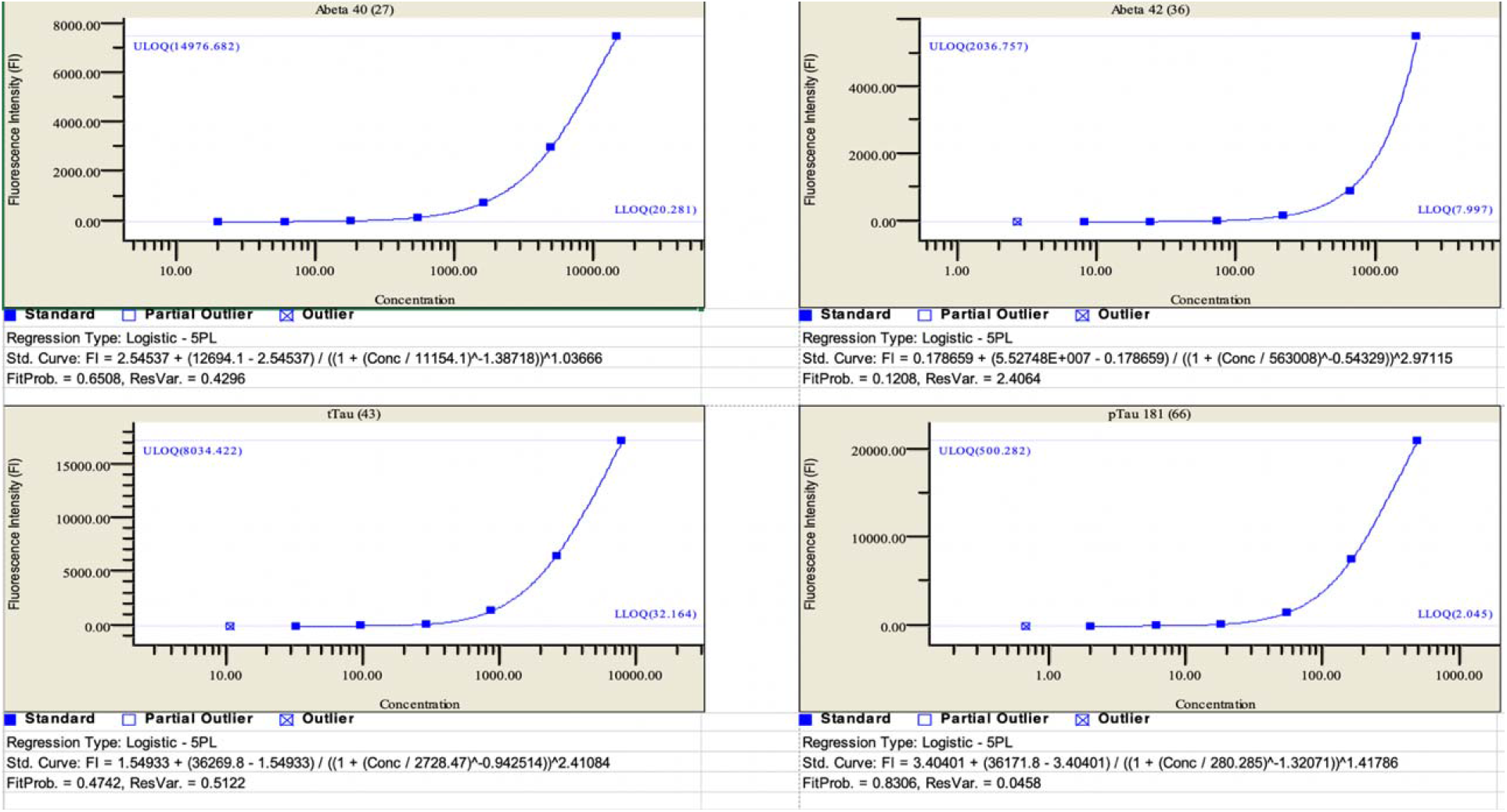
LLQQ levels of CSF proteins.

**Supplementary Table 1:** List of antibodies used.

| Antibody | Target | Identifier |
| --- | --- | --- |
| Mouse anti-Aβ (McSA1) | Aβ38-42 | MM-0015-1P |
| Rabbit anti-human Aβ | Aβ42 | JP28051 |
| Mouse anti-Iba1 | Ionized calcium binding adaptor molecule 1 (Iba1) | ab15690 |
| Rabbit anti-oligomer (A11) | Aβ42 oligomers | AHB0052 |
| Rabbit anti-amyloid fibrils (OC) | Generic epitopes common to amyloid fibrils and fibrillar oligomers | AB2286 |
| Mouse anti-MC1 | AD-specific conformational modification of tau | Peter Davies, Department of Pathology, Albert Einstein College of Medicine |
| Rabbit anti-TREM2 | TREM2 receptor | MA530971 |
| Rabbit anti-MAP2 | Microtubule-associated protein 2 | ab183830 |
| Rabbit anti-LAMP1 | Lysosomal associated membrane protein 1 | L1418 |
| AT8 | Tau phosphorylated at serine 202 and threonine 205 | MN1000 |
| Mouse anti-HT7 | MAPT | MN1000 |

